# Genome-Wide SNPs Reveal the Drivers of Gene Flow In An Urban Population of the Asian Tiger Mosquito, *Aedes albopictus*

**DOI:** 10.1101/173963

**Authors:** Thomas L. Schmidt, Gordana Rašić, Dongjing Zhang, Xiaoying Zheng, Zhiyong Xi, Ary A. Hoffmann

## Abstract

*Aedes albopictus* is a highly invasive disease vector with an expanding worldwide distribution. Genetic assays using low to medium resolution markers have found little evidence of spatial genetic structure even at broad geographic scales, suggesting frequent passive movement along human transportation networks. Here we analysed genetic structure of *Ae. albopictus* collected from 12 sample sites in Guangzhou, China, using thousands of genome-wide single nucleotide polymorphisms (SNPs). We found evidence for passive gene flow, with distance from shipping terminals being the strongest predictor of genetic distance among mosquitoes. As further evidence of passive dispersal, we found multiple pairs of full-siblings distributed between two sample sites 3.7 km apart. After accounting for geographical variability, we also found evidence for isolation by distance, previously undetectable in *Ae. albopictus*. These findings demonstrate how large SNP datasets and spatially-explicit hypothesis testing can be used to decipher processes at finer geographic scales than formerly possible. Our approach can be used to help predict new invasion pathways of *Ae. albopictus* and to refine strategies for vector control that involve the transformation or suppression of mosquito populations.

**Author Summary:** *Aedes albopictus*, the Asian Tiger Mosquito, is a highly invasive disease vector with a growing global distribution. Designing strategies to prevent invasion and to control *Ae. albopictus* populations in invaded regions requires knowledge of how *Ae. albopictus* disperses. Studies comparing *Ae. albopictus* populations have found little evidence of genetic structure even between distant populations, suggesting that dispersal along human transportation networks is common. However, a more specific understanding of dispersal processes has been unavailable due to an absence of studies using high-resolution genetic markers. Here we present a study using high-resolution markers, which investigates genetic structure among 152 *Ae. albopictus* from Guangzhou, China. We found that human transportation networks, particularly shipping terminals, had an influence on genetic structure. We also found genetic distance was correlated with geographical distance, the first such observation in this species. This study demonstrates how high-resolution markers can be used to investigate ecological processes that may otherwise escape detection. We conclude that strategies for controlling *Ae. albopictus* will have to consider both passive reinvasion along human transportation networks and active reinvasion from neighbouring regions.

## Introduction

The Asian Tiger mosquito, *Aedes albopictus,* is one of the world’s most dangerous invasive species (Global Invasive Species Database, http://www.issg.org/database/). It has been implicated in recent outbreaks of Chikungunya [1] and Dengue [2] in the tropics and even temperate regions [3, 4], and its hyperaggressive diurnal bloodfeeding reduces the quality of human environments [5]. Minimising the negative impact of *Ae. albopictus* with pesticides has proven increasingly difficult due to insecticide resistance, particularly among populations within the native distribution of this species [6-9], which extends from India to South East Asia to Japan [10].

Modification of mosquito populations using the endosymbiotic bacterium *Wolbachia* has been proposed as a viable alternative to pesticide-based population suppression [11]. This study considers an urban population of *Ae. albopictus* in Guangzhou, China, that has undergone releases of *Wolbachia-*infected males to suppress populations at two locations since March 2015. Guangzhou region is within the native distribution of *Ae. albopictus*, and experiences occasional Dengue outbreaks despite local absence of the primary vector of Dengue, *Ae. aegypti* [9]. *Aedes albopictus* females in Guangzhou are normally superinfected with two *Wolbachia* strains, *w*AlbA and *w*AlbB [12], and *Wolbachia-*based population suppression has been conducted by releasing males that carry an additional strain, *w*Pip, that has been transferred from *Culex pipiens* (Diptera: Culicidae) [13]. This population suppression effort has been successful, with population reductions of more than 90% maintained over two years (Xi pers. comm.). However, whether *Wolbachia* releases are able to suppress *Ae. albopictus* populations sustainably over a large geographic region depends on the frequency at which suppressed sites are reinvaded by mosquitoes dispersing into the sites from outside. This reinvasion determines the frequency and intensity at which releases need to be repeated to maintain population suppression.

To estimate the susceptibility of a region to reinvasion an understanding of basic dispersal parameters in *Ae. albopictus* is needed. Dispersal in *Ae. albopictus* is generally thought to represent a combination of short-range active flight of adults and long-range passive movement of immature stages along human transportation networks [10, 14-17]. The success of *Ae. albopictus* as an invasive species likely reflects its ability to disperse along human transportation networks such as shipping routes [10] and roads [14]. This has contributed to little genetic differentiation at broad scales throughout its range [18-21], with most genetic variation observed within sampling sites [15, 22]. This pattern of genetic structuring is typical for newly colonised populations experiencing founder effects, but *Ae. albopictus* populations show little differentiation even within their native range [16, 17, 23, 24]. Some studies in recently invaded regions have reported mild genetic structure [22, 25], though the spatial dependence of this has not been tested.

Although negligible genetic differentiation has been observed between *Ae. albopictus* samples within countries [17, 18] and cities [26], structure at these scales is regularly observed in *Ae. aegypti*, the primary Dengue vector [27-31]. This seems incongruous given that the estimates of the flight range potential in *Ae. albopictus* are comparable to those in *Ae. aegypti* (100m - 500m: [10, 32-36]. At finer scales we expect active dispersal to have a stronger influence on genetic structure and contribute to patterns of spatial genetic structure such as isolation by distance (IBD [37]). However, at broader scales that exceed species’ active dispersal capacity, a relationship between spatial distance and genetic distance such as IBD weakens and genetic structure is driven more by patterns of passive dispersal such as via trade or traffic routes. Because strong genetic structure among *Ae. albopictus* populations has yet to be observed at any scale, passive dispersal has been proposed as the most important determinant of genetic structure in this mosquito [38].

The lack of observed genetic differentiation at finer spatial scales may also be an artefact of using genetic markers that lack the power to detect subtle genetic patterns [15]. Analyses of *Ae. aegypti* at genome-wide single nucleotide polymorphisms (SNPs) discovered through double-digest restriction-site associated DNA sequencing (ddRADseq [39]) have provided a wealth of new ecological information at fine scales [29-31, 40, 41], including direct evidence of passive dispersal and spatial genetic structure at distances < 4km [31]. Until now, genetic patterns in *Ae. albopictus* have been studied using allozymes, mtDNA, and microsatellites ([17-21]; see review in Goubert et al. [15]), that have a much lower resolution than genome-wide SNPs in deciphering genetic patterns in *Ae. aegypti* [31, 41].

In this study, we used ddRAD sequencing to genotype *Ae. albopictus* from Guangzhou, China at genome-wide SNPs to test explicit hypotheses about the processes shaping the genetic structure within the city. Specifically, we determined the relative effects on genetic structure of IBD, passive dispersal along human transport networks, and barriers to dispersal. We sampled *Ae. albopictus* in sites closely situated among networks of human transportation to test hypotheses of the contribution of active and passive dispersal on genetic structure. We also investigated whether human transportation networks such as highways or rivers could act as barriers to dispersal. Highways have been observed to restrict dispersal in *Ae. aegypti* [31, 42, 43], and given that *Ae. albopictus* tends to prefer sylvan, rural and suburban habitats over urban ones [10, 44], large urban highways such as those in Guangzhou may pose particularly effective barriers to mosquito dispersal. There is considerable evidence for passive dispersal along human transport networks and no evidence of IBD within countries [14, 15, 26, 45, 46], and our study is the first to employ high-density genomic markers to investigate gene flow within a city. This paper describes how with these markers we were able to find evidence for both IBD and passive dispersal along human transportation networks, but no evidence for dispersal barriers.

## Methods

### Sampling

*Aedes albopictus* were collected from 12 sites on public land in Guangzhou, China, between September 23^rd^ and October 22^nd^, 2015. The sample sites were distributed across an area of 380 km² through 9 administrative districts, and were separated by distances ranging from 2.73 km to 49.34 km (Fig 1). All subsequent references to sites follow the numerical designations in Fig 1. At each site, natural containers were searched for larvae and pupae that were collected and then combined into a single sample to be reared in the laboratory until eclosion.

**Fig 1:**
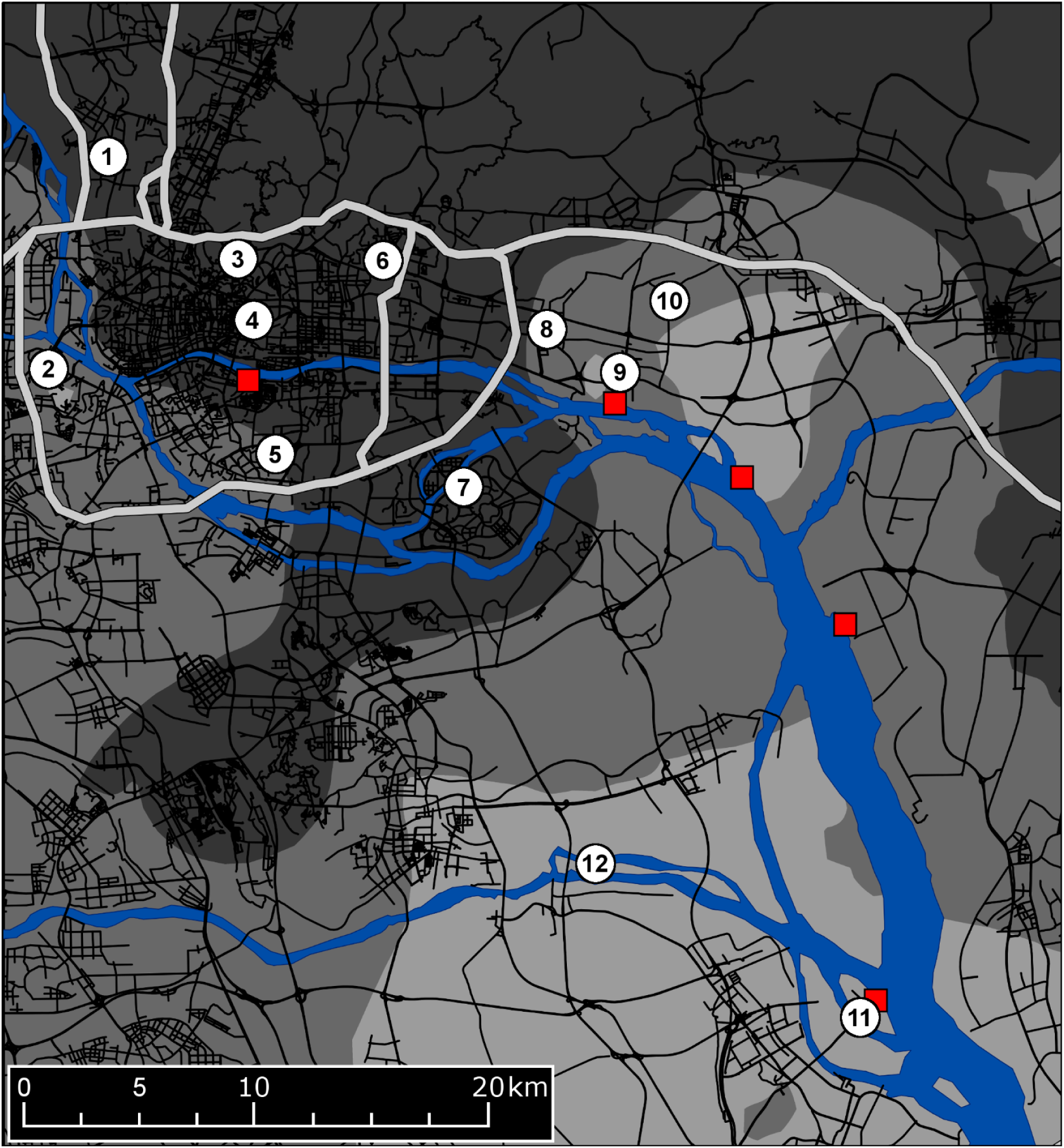
Sampling locations and study site geography. The numbered yellow circles show the 12 sample sites. The red squares show the locations of the five shipping terminals. The rivers and estuary are shown in blue. High-capacity highways are plotted as wide, light grey lines, while the greater highway network is in black. The background shading describes the average interpolated Breteau Index score: 0-2 (dark grey); 2-4 (grey); 4-6 (light grey). The underlying road network is derived from the Chinese “Highways” shapefile made available by MapCruzin.com and OpenStreetMap.org under the Open Database License [https://opendatacommons.org/licenses/odbl/1.0/]

Samples from Sites 1 – 8 were raised in tap water containing shrimp powder, while samples from Sites 9 – 12 were raised in tap water containing shrimp powder, bovine liver powder and nutritional yeast. Following eclosion, adults were fed on 10% sugar solution and allowed to mate, but not bloodfeed. After 4 - 6 days, 20 females from each sample were killed by freezing and stored in ethanol until DNA extraction. Genomic DNA was extracted using Roche DNA Isolation Kit for Cells and Tissues (Roche, Pleasanton, CA, USA), with an additional step of RNAse treatment. From the 12 sample sites, we selected 152 individuals for ddRAD sequencing.

### SNP discovery

#### *In silico* restriction enzyme selection

We used the program *DDsilico* [41] to perform *in silico* digestions of the *Ae. albopictus* genome assembly AaloF1 [47] using different restriction enzyme pairs. *DDsilico* calculates how many sequenceable fragments of a given bin size can be produced for each enzyme pair. Two frequent cutter enzymes, *MluC*I and *Nla*III, produced an optimal number of fragments for a library size of 50 within a fragment length range 350 - 450bp and desired average read depth > 20. This enzyme combination has also been used to construct ddRADseq libraries in *Ae. aegypti* [29-31, 41].

#### Double-digest RADseq library preparation

We performed an initial digestion of 100 ng of genomic DNA in a 40 µL reaction, using 10 units each of *MluC*I and *Nla*III restriction enzymes (New England Biolabs, Beverly MA, USA), NEB CutSmart^®^ buffer, and water. Digestions were run for 3 hours at 37°C with no heat kill step, and the products were cleaned with 60 µL Ampure XP™ paramagnetic beads (Beckman Coulter, Brea, CA). These were ligated to modified Illumina P1 and P2 adapters overnight at 16°C with 1000 units of T4 ligase (New England Biolabs, Beverly, MA, USA), followed by a 10-minute heat-deactivation step at 65°C.

We performed size selection with a Pippin-Prep 2% gel cassette (Sage Sciences, Beverly, MA) to retain DNA fragments of 350 - 450 bp. Final libraries were created by pooling eight 10 μL PCR reactions per library, each consisting of 1 μL size-selected DNA, 5μL of Phusion High Fidelity 2× Master mix (New England Biolabs, Beverly MA, USA) and 2 μL of 10 μM standard Illumina P1 and P2 primers, run for 12 PCR cycles. These were cleaned and concentrated using an 0.8× concentration of Ampure XP™ paramagnetic beads (Beckman Coulter, Brea, CA) to make the final libraries. Three libraries containing a total of 152 *Ae. albopictus* were sequenced in three Illumina HiSeq2500 lanes using 100 bp paired-end chemistry.

#### Data processing and genotyping

We processed raw fastq sequences within the customized pipeline of Rašić et al. [41], with reads filtered with a minimum phred score of 13 and trimmed to the same length of 80 bp. High-quality reads were aligned sequentially to the *Ae. albopictus* reference nuclear genome AaloF1 [47] followed by the *Ae. albopictus* mitochondrial genome, and then the remaining unaligned sequences were aligned concurrently to the genomes of the *Wolbachia* strains *w*AlbB [48] and *w*Pip [49], with all alignments performed with the program Bowtie [50]. We allowed for up to 3 mismatches in the alignment seed, and uniquely aligned reads were analysed with the Stacks pipeline [51], which we used to call genotypes at RAD stacks of a minimum depth of 5 reads. We observed an average depth of 23.33 reads among the AaloF1 alignments retained after quality filtering.

The alignment to *w*Pip was necessary as one of the sampling sites (Site 11) had undergone releases from March 2015 of *Ae. albopictus* males carrying this *Wolbachia* strain in addition to the naturally occurring *w*AlbA and *w*AlbB. Although mosquitoes were irradiated before release to ensure any females remaining in the sample could not reproduce and introduce the *w*Pip strain into the population [52], there was a slight possibility that accidental introduction could have occurred. We therefore tested if Shazi Island mosquitoes showed higher rates of alignment to the *w*Pip genome than other study sites. Dadaosha Island (Site 12) also underwent *w*Pip male-releases, but these began in March 2016 after the sampling for our study had taken place.

The Stacks program Populations was used to export VCF files for additional filtering with the program VCFtools [53]. We first removed 9 individuals with missing data > 20%, then applied further filtering to retain only those loci that were at Hardy-Weinberg Equilibrium, present in ≥ 75 % of individuals and with minor allele frequencies of ≥ 0.05. To avoid using markers in high linkage disequilibrium, data were thinned so that no single SNP was within 250 kbp of another. As the *Aedes* genome is thought to contain approximately 2.1 megabases per cM [54], 250 kbp roughly corresponds to eight SNPs per map unit, a sampling density that has been shown to largely eradicate the effects of linkage in SNPs [55]. Our final dataset had 3084 unlinked and informative SNPs for analyses of relatedness and genetic structure.

### Relatedness analysis

Analyses of population structure are generally biased when closely-related individuals such as full-siblings are included in the analyses [56]. To identify full-sibling relationships among our samples, we calculated Loiselle’s *k* [57] using the program SPAGeDi [58]. First and second-degree kin relations can be ascertained with confidence using large SNP datasets [31, 39, 59-61], wherein individuals with pairwise *k* > 0.1875 are full-siblings, and those with 0.1875 > *k* > 0.09375 are half-siblings, if each pair is assigned the most likely kinship category [62]. We sampled one individual from each putative full-sibling group with the smallest percentage of missing data, leaving 116 individuals for analysis of genetic structure.

We were also interested in observing related pairs at different sampling sites. Larval *Ae. aegypti* full-siblings have been caught in traps up to 1.3 km apart [31], well above the active dispersal range of *Ae. albopictus* [10, 32-35]. As the minimum distance between sites in this study was more than double this distance, movement at these scales would most likely also be human-mediated. To avoid incorrectly identifying the relationship of closely-related pairs we used the program ML-Relate [63] to perform specific hypothesis tests of relationship. For each pair we ran one test that estimated the relationship assuming that the kinship category assigned using *k* was more likely than the next most likely kinship category, followed by tests that assumed that the kinship category assigned using *k* was less likely to be correct. Thus, for pairs with *k* > 0.1875, tests would determine whether the pair were full-siblings or half-siblings, while for pairs with 0.1875 > *k* > 0.09375 tests would help determine whether the pair were full-siblings, half-siblings or unrelated. Tests were run using 10,000,000 simulations of random genotype pairs for each.

### Analysis of genetic structure

#### Site geography

We first identified local geographical features that could potentially act as either dispersal barriers or as conduits for passive dispersal. Potential dispersal barriers were high capacity highways and major river tributaries and conduits for passive dispersal were rivers, highways and ports/shipping terminals. High capacity highways were designated following the key of the “World Street Map” basemap file in ArcMap 10.3.1 from ESRI [64]. Major river tributaries included the Zhujiang and East Rivers, the Shawan and Lianhuashan waterways, and the Shizi Ocean estuary. To derive a continuous variable representing each barrier type, individuals from a given sampling site were assigned an integer from 0 - 3 that described their position relative to barriers. For highways (“Highway Barriers”) these positions were: Northern group (0; Site 1), Central group (1; Sites 2-6) and Southern group (2; Sites 7-12). For rivers (“River Barriers”) the positions were Northern group (0; Sites 1, 3, 4, 6, 8, 9 and 10), Central group (1; Sites 5 and 7), Southern group (2; Site 2) and Southernmost group (3; Sites 11 and 12) (Fig 1).

To investigate passive dispersal along the river, highway and port networks we first generated a score for each individual for each of the three network types describing its isolation from the network. Isolation from the river network (“River Isolation”) was calculated as the log-transformed Euclidean distance to the nearest river segment, while isolation from the port network (“Port Isolation”) used the distance to the nearest of five shipping terminals (Henan Terminal, Huangpu Old Port, Huangpu New Port, Xinsha Port, and the Shazi Island Automotive Terminal). We assessed isolation from highway network (“Highway Isolation”) by constructing a metric as the square root of the total area within a 1 km radius not covered by the highway network, assuming an average highway width of 30 m. A similar metric calculated for the railway network was omitted due to high correlation with the highway network (R² = 0.695, P < 0.001). The highway network was constructed based on the “Highways” shapefile from MapCruzin (http://www.mapcruzin.com/free-china-country-city-place-gis-shapefiles.htm). Positions of sample sites relative to each network are shown in Fig 1.

To analyse the influence of each network on individual genetic distances, we constructed pairwise additive matrices by summing the two scores of each individual pair to create a total isolation score. Individuals from the same sample site were assigned a total isolation score of zero. For all analyses of network influence, we considered only network connectivity and not distance travelled along the network, as this would require the estimation of the probability distribution of dispersal distances along networks, for which no data was available. Instead of treating each network as a resistance surface with frequency of dispersal declining with distance, our method effectively modelled dispersal along each network with a continuous uniform distribution.

#### Network habitat quality

The null expectation is that individuals from sites scoring low for network isolation would have lower than average genetic distances, as these highly-connected sites will act as “hubs” for regional gene flow. However, for these sites to operate in this way they must also be situated in areas of suitable habitat, so that immigrants do not face high mortality or a lack of oviposition sites after a dispersal event, and so that there will be a large number of migrants (*Nm* [65]) dispersing from these sites to others across the network. For this reason, we investigated the relationships between each network variable and two variables representing habitat quality: the Breteau Index (number of containers with larvae or pupae per 100 dwellings) and the Normalized Difference Vegetation Index (NDVI), given that *Ae. albopictus* prefers sylvan or rural environments in its native range [44].

We calculated the Breteau Index from fortnightly data available on the Guangzhou Centre for Disease Control and Prevention website (http://www.gzcdc.org.cn/). We averaged Breteau Indices from 85 locations over the three months prior to sampling, then interpolated the scores using Ordinary Kriging in ArcMap 10.3.1 [64]. Kriging was performed with an exponential semivariogram model and a 36-point nearest-neighbour search function. The Kriging results are shown in Fig 1. The derived raster layer was used to calculate an average Breteau Index within a 1 km radius for each individual. We also calculated the average NDVI within a 1 km radius for each individual, using a NDVI raster from ArcGIS Online (http://www.arcgis.com/home/item.html?id=2354af91ee294bd3825c27dfa914a9a7), assigning values from -1 (no vegetation) to 1 (completely vegetated). The two estimates of habitat quality were positively correlated, but not at 95% confidence (R² = 0.298, P < 0.066).

#### Spatial genetic structure

We used SPAGeDi to calculate Nei’s *D* [66] as a measure of group genetic distance between sample sites, and Rousset’s *a* [67] for a measure of genetic distance between individuals. We performed an AMOVA [68] with the function *amova* (R package “pegas” [69] to partition the variance observed within and among sample sites. All other operations were performed with the package “vegan” [70]. We performed Mantel tests [71], partial Mantel tests [72] and distance-based redundancy analysis [73] to assess relationship between the landscape variables and genetic distance. We first used Mantel tests (function *mantel*) to investigate correlations between genetic distance, geographical distance and each of the five landscape variables. We then used paired Mantel tests (function *mantel.paired*) to investigate the contribution of each variable with the effects of geographic distance being removed, and *vice versa* (testing the effect of geographic distance after removing each landscape variable).

Before constructing dbRDA models, we first used the function *pcnm* to transform the matrices of geographical distance and additive network isolation into principal components (PCs). We built two dbRDA models: the first treating geographic distance as an explanatory variable (dbRDA1), and the second where geographic distance effects were partialled out (dbRDA2). We constructed dbRDA1 using the first two PCs from each of the distance and network matrices. In dbRDA2, we performed an initial dbRDA of the PCs of distance on genetic distance and retained the four PCs that were significant (P < 0.05 following Benjamini-Hochberg correction [74], which were placed in the conditional matrix. We then used the first two PCs from each of the network matrices for model construction.

Terms were added to each model sequentially using the adjusted R² as a measure of model fit, and construction was finished once adjusted R² reached a maximum. Marginal significance of terms in the final models was assessed using an ANOVA, with P-values adjusted using the Benjamini-Hochberg correction. We used Eta squared (η^2^ [75]) as a measure of effect size. The functions *capscale*, *anova.cca* and *p.adjust* (package “stats”) were used for dbRDA, significance testing with ANOVA and P-value adjustment, respectively.

#### Type I error testing

Ordination methods such as dbRDA, and to lesser extent partial Mantel tests, are known to suffer from inflated Type I error [76]. Specifically, Type I errors can be caused if hypotheses of landscape influence are tested only against the null hypotheses of panmixia, rather than against alternative geographical hypotheses such as IBD [77, 78]. A Mantel test between log-transformed geographical distance and Rousset’s *a* showed significant spatial autocorrelation potentially indicating IBD (r = 0.062, P < 0.001, Table 1). When genetic structure exhibits IBD, spatial dependence among landscape variables and inadequately extensive sampling can lead to significance being observed when no landscape effect exists [79]. We performed further analyses that would identify if any of our analyses were at high risk of Type I error.

We adapted the method of Kierepka and Latch [79] to test for Type I error inflation in our partial Mantel tests and final dbRDA models. We used CDPOP [80] to simulate the field site without any geographical features, creating an artificial environment in which only IBD could explain the pattern of structuring. We designated 1500 locations as the positions of individual mosquitoes, which included the locations of the 116 individuals used in genetic structure analyses and 1384 new locations placed throughout the field site. New locations were placed at a density corresponding to the interpolated Breteau Index scores, so that regions scoring 2-4 had three times the density of individuals as those scoring 0-2, and regions scoring 4-6 had five times the density (see Fig 1). As CDPOP does not allow individuals to occupy the same space, we added “noise” to the locations of the 116 real individuals, moving each individual up to 250m in a random location, which was less than the median dispersal distance. We used parameters of high recruitment and high mortality, and negative exponential dispersal approximating a leptokurtic dispersal kernel with no maximum limit to dispersal distance, but where most dispersal took place within 500 m. The CDPOP input file listing all relevant parameters is supplied in S1 Table.

We allowed CDPOP to generate genotypes for each individual, so that the population would initially be at maximum genetic diversity. To ensure the real and simulated data sets would have similar numbers of effective alleles, we used fewer SNPs in our simulations. Effective allele counts have been found to correspond well with analytical power across different types of genetic data [81, 82]. We used the method of Kimura and Crow [83] to calculate the number of effective alleles in our 3084 loci set, which gave 3694 effective alleles, corresponding to 1847 biallelic loci at maximum diversity.

We constructed 100 simulations and ran each for 100 discrete generations, after which we sampled the genotypes of the mosquitoes present at the original 116 locations. We used Mantel tests to test if all simulation samples showed similar levels of IBD (r, x̅ = 0.059 ± 0.019, all P < 0.05) as in the empirical data (r = 0.062, P < 0.05). For each simulation sample we constructed new partial Mantel and dbRDA models containing the same parameters as the observed data, and tested for the marginal significance of each variable. As significance in our models was assessed at 95% confidence, we considered that if more than 5 of the 100 simulated samples showed significance for a variable then observations regarding that variable would be at elevated risk of Type I error.

## Results

### Relatedness analysis

Comparisons across sample sites revealed four probable full-sibling kinship groups (0.203 ≤ *k* ≤ 0.474, 0.10% of pairs) and a single probable half-sibling group (*k* = 0.156; 0.02% of pairs) with members spread between two sample sites 3.657 km apart (Fig 2), Jiuwei Village and Zhucun Village (Sites 8 and 9). This distance is much greater than flight range estimates of *Ae. albopictus* [32-35], which suggests that this dispersal likely reflects passive transportation by humans. Specific tests of relationship using maximum-likelihood estimation found that, of the nine pairs of (*k* > 0.1875) found across sites, six were definitely full-siblings and not half-siblings (P < 0.001), while the remaining three could be of either category.

**Fig 2:**
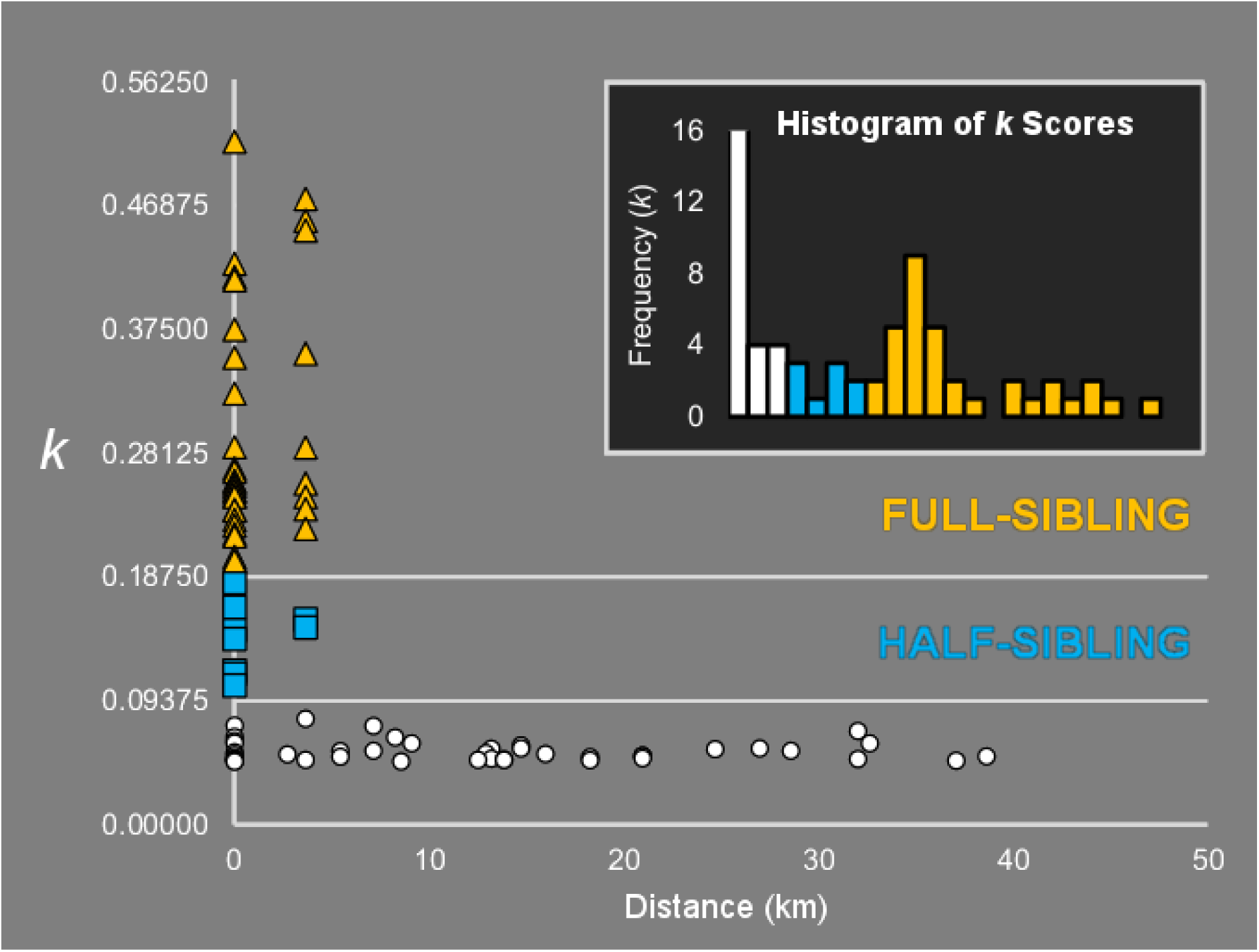
Loiselle’s *k* scores for pairs of individuals and (inset) a histogram showing the frequencies of these scores. Most related pairs were found within sample sites, but several pairs were found between two sites 3.657m apart. The half-sibling pair with the lowest *k* showed clear separation from all pairs of lower relatedness. Few related pairs had *k* scores close to the full-sibling/half-sibling category boundary (inset).

Out of 854 comparisons between individuals sampled within the same site, 25 pairs (2.93%) were full-siblings (0.200 ≤ *k* ≤ 0.517) and 7 pairs (0.82%) were half-siblings (0.105 ≤ *k* ≤ 0.183). A histogram of *k* scores showed fewer scores near *k* = 0.1875, the full-sibling/half-sibling category boundary, with the number of full-sibling pairs increasing sharply with *k* (Fig 2, inset). We did not observe the same pattern at the half-sibling/unrelated category boundary, but the most distantly related half-sibling pair had a *k* score 31.2 % larger than the most genetically similar pair of unrelated individuals.

### Spatial genetic structure

The AMOVA showed significant genetic structuring among sampling sites (P < 0.001), accounting for 8.1% of the observed variation (η² = 0.081). A simple Mantel test on matrices of individual genetic distance and log-transformed geographic distance showed a small but significant correlation (Table 1, r = 0.062, P < 0.001). Mantel tests for other landscape variables showed that “Port Isolation”, “River Isolation” and “Highway Barriers” were all better predictors of genetic distance than geographical distance. (Table 1). Neither “Port Isolation” nor “River Isolation” showed any correlation with distance, while the other variables all did.

**Table 1:** Results of Mantel and partial Mantel tests.

|  | *Mantel Tests* |  | *Partial Mantel Tests* |  |
| --- | --- | --- | --- | --- |
|  | *a* | Distance | *a* \| Distance | Type I error |
| Port Isolation | 0.077 | -0.029 | 0.070 | 2 |
| River Isolation | 0.075 | -0.359 | 0.047 | 6 |
| Highway | 0.052 | 0.432 | -0.041 | 0 |
| Highway Barriers | 0.150 | 0.459 | 0.137 | 21 |
| River Barriers | -0.051 | 0.530 | -0.082 | x |
| Distance | 0.062 | . | . | . |
| | | $P < 0.05$ | $P < 0.01$ | $P < 0.001$ |

Cells contain $r$ scores and are shaded by degree of statistical significance. Mantel tests are between landscape variables (including geographic distance) and Rousset's $a$ , and between landscape variables and geographic distance. Partial Mantel tests are between landscape variables and Rousset's $a$ , after accounting for geographic distance. The number of simulations (out of 100) showing significance is indicated for each partial Mantel test.

#### Isolation-by-geographic distance

Nei’s *D* scores between all sample sites showed a weak but significant positive correlation with distance (R² = 0.088, P < 0.016), with considerable variability at small spatial scales. However, these broad-scale analyses could be potentially biased by the two sites in the south, Dadaosha Island (Site 11) and Shazi Island (Site 12), as in addition to them being distant from the northern sites they were also semi-rural as opposed to urban and scored more highly for “Highway Isolation” (7.98 ± 3.47) than the other sites (2.72 ± 1.15). Therefore, for an analysis containing all pairwise comparisons, we could not separate the effects of spatial and non-spatial influences on genetic structure.

To explore this further, we investigated the spatial dependence of pairwise Nei’s *D* scores with several subsets of sites, as detailed in Fig 3. We first identified one sampling site at the North Campus of Sun Yat-sen University (Site 4; red triangles) as highly differentiated from other sites, with Nei’s *D* scores 75.8% higher than the average for all other sites. We excluded this site from analyses, leaving the two southern sites (Sites 11 and 12) and the nine northern sites (Sites 1, 2, 3, 5, 6, 7, 8, 9 and 10). As “Highway Isolation” showed no spatial dependence among the northern sites (R² = 0.003, P < 0.751), we could investigate patterns of IBD among these sites which would not be confounded by the highway network. We first analysed the nine northern sites only (white crosses on grey), which showed mild IBD (R^2^ = 0.136, P < 0.027) at distances of 3.66 - 26.98 km. We then performed separate analyses between the northern sites and Site 11 (blue squares) and Site 12 (yellow diamonds), which showed strong IBD at distances of 17.23-37.08 km (R^2^ = 0.441, P < 0.050) and 28.57-49.34 km (R^2^ = 0.540, P < 0.024) respectively. These results suggest a pattern of IBD that is unconfounded with the effects of highway, port or river networks.

**Fig 3:**
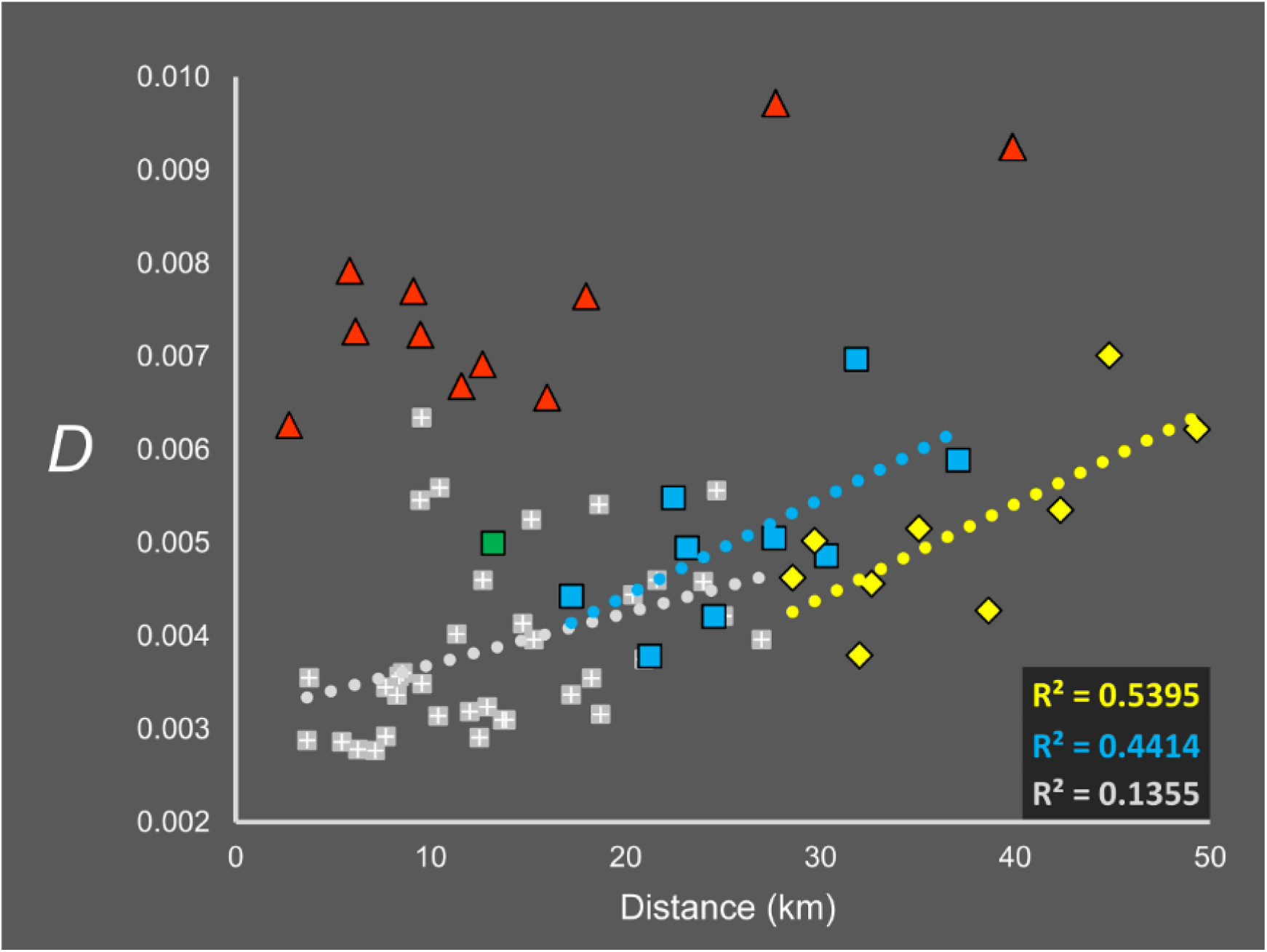
Relationship between pairwise Nei’s *D* estimates and geographic distance. Red triangles show distances for Site 4, which was highly differentiated from all others. Yellow diamonds denote distances between Sites 1,2,3,5-10 and Site 11, blue squares denote the same but for Site 12. Distances between Sites 11 and 12 are shown by the green square. White on grey crosses show distances for Sites 1,2,3,5-10.

#### Isolation-by-landscape distance

Partial Mantel tests on “Port Isolation”, “River Isolation” and “Highway Barriers” were all significant after removing the effects of geographic distance (Table 1). However, simulations showed that both the “River Isolation” and “Highway Barriers” models, had an inflated Type I error. Thus, we considered the effects of “Highway Barriers” and “River Barriers” on genetic structure to be untestable with our sampling design.

The dbRDA1 model with all variables showed that the “Port Isolation” contributed the most to genetic structure, followed by the effect of geographic distance (Table 2). However, once we calculated Variance Inflation Factors with the function *vif.cca,* we found high collinearity between the variables. Construction of dbRDA2 excluded the “River Barriers” variable, and again, the “Port Isolation” variable was the most important in the model. Variance Inflation Factors were less than 5% between the independent variables. “Highway Barriers” contributed the least to the model and showed a much smaller effect size than the three network isolation variables. None of the variables were significant in > 5 % of simulations i.e. they did not show evidence for an inflated Type I error.

**Table 2:** Results of dbRDA1 and dbRDA2.

|  | *dbRDA1* | *dbRDA2* |  |  |
| --- | --- | --- | --- | --- |
|  | Order | Order | Effect size | Type I error |
| Port Isolation | 1st | 1st | 0.0265 | 5 |
| River Isolation | 3rd | 2nd | 0.0248 | 5 |
| Highway Isolation | 6th | 3rd | 0.0238 | 5 |
| Highway Barriers | 4th | 4th | 0.0145 | 5 |
| River Barriers | 5th | . | . | . |
| Distance | 2nd | Conditional |  |  |
|  |  | P < 0.05 | P < 0.01 | P < 0.001 |

The order of inclusion of variables is indicated for each model. Cells containing the effect sizes of variables retained in dbRDA2 are shaded by statistical significance. The number of simulations (out of 100) showing significance is indicated for each variable in dbRDA2.

### mtDNA haplotype and *Wolbachia* strain distributions

Mitochondrial DNA haplotype variants were broadly distributed throughout the study area (Fig 4). A single common haplotype was found at all 12 sites and two others were found at seven and three sites respectively. No other haplotypes were found at more than one location. Haplotype variants were separated from the common haplotype by 4 polymorphic sites at most, representing a divergence of 0.024%. Analyses demonstrating an overall lack of spatial clustering among mitochondrial haplotypes are described in S2 Table and S1 Fig.

**Fig 4:**
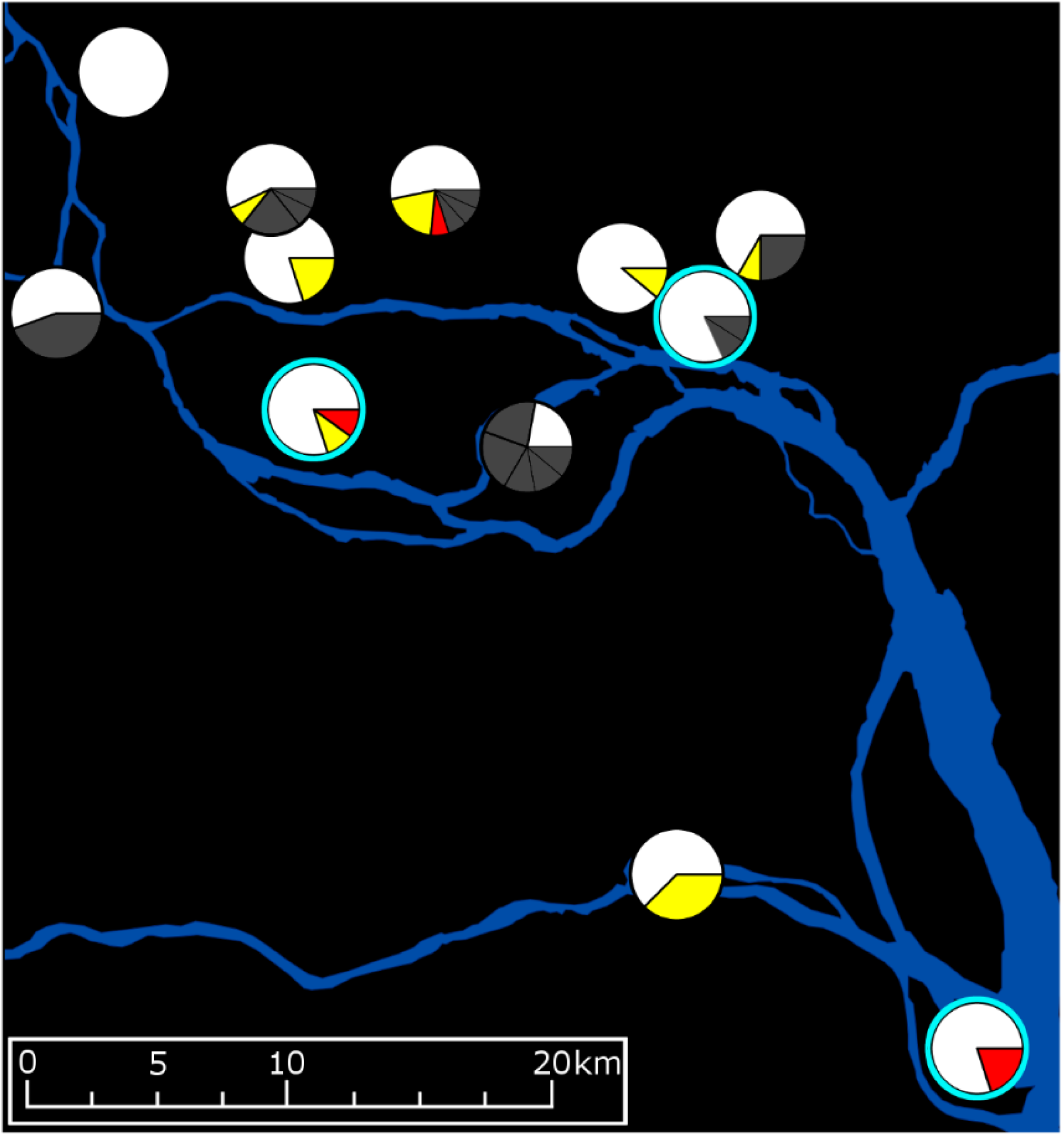
Locations of mtDNA and *w*AlbB haplotype variants. Pie charts represent mtDNA haplotypes, with the following legend: white = the single common haplotype; yellow = rare haplotype A, 1 SNP; red = rare haplotype B, 3 SNPs; dark grey = rare haplotypes of variable divergence, each confined to one sample site. One SNP in *w*AlbB was found in three individuals from different sites, these are indicated with a blue ring.

Among *w*AlbB haplotypes we identified 3 SNPs that were found in more than one individual. One of these was observed in three individuals from different sites (Sites 5, 9 and 11; see Figs 1 & 4). Comparing the ratios of *w*Pip to *w*AlbB alignments among the Shazi Island individuals with all individuals, there was no statistically higher rate of alignment to *w*Pip in Shazi Island (x̅ = 0.489 ± 0.037) than the average (x̅ = 0.479 ± 0.083), so we considered the sample not to be biased by the releases of males with the *w*Pip infection.

### Network habitat quality

The Breteau Index was strongly and positively correlated with “Highway Isolation” (R² = 0.641, P < 0.002). “River Isolation” (R² = 0.291, P < 0.071) had a negative but nonsignificant correlation with the Breteau Index and “Port Isolation” (R² = 0.002, P < 0.900) showed no correlation. The same general pattern was observed between the NDVI and “Highway Isolation” (R² = 0.300, P < 0.066), “River Isolation” (R² = 0.072, P < 0.400) and “Port Isolation” (R² = 0.019, P < 0.666), though none of the relationships were significant at the 95% confidence level. These results suggest that as connectivity to the highway network increases there is a corresponding decrease in *Ae. albopictus* habitat quality.

## Discussion

Our results show spatially explicit genetic structure in *Ae. albopictus* within a city in its native range, revealed with high resolution markers. Additionally, our hypothesis testing around variables that contribute to the observed genetic structure shows that this structure likely reflects a complex interaction between various forms of active and passive dispersal, and is broadly consistent with expectations of occasional long distance gene flow and local genetic drift [15-17]. We found evidence for IBD and gene flow along human transportation networks, with proximity to ports being the strongest predictor of genetic distance in all analyses. Further evidence of passive dispersal came from the discovery of several pairs of closely related individuals at sample sites > 3.6 km apart and from the broad spatial distribution of rare mtDNA and *Wolbachia* haplotype variants. We found no evidence for geographical barriers to gene flow at this spatial scale. As transportation networks and geographical distance both showed similar power in predicting genetic distances, we propose that passive gene flow and IBD can operate at similar intensities at this spatial scale.

The structure observed between populations in this study (8.1 %) is comparable with [22, 25] or higher than [26] findings from some spatially unexplicit studies at similar scales. However, identifying IBD patterns in *Ae. albopictus* has proven difficult with a range of molecular markers [17-21] that have lacked the power to investigate population processes at fine scales [18, 26]. Our analysis of city-wide genetic structure using genome-wide SNPS revealed a pattern of IBD among *Ae. albopictus* in Guangzhou. Specifically, among the northern sites (Sites 1, 2, 3, 5 - 10), genetic distances increased with spatial distances irrespective of whether the northern sites were analysed among themselves or whether they were analysed in relation to either of the two southern sites (Sites 11 and 12). As these northern sites were all highly urbanised and their isolation scores from each of the highway, river and port networks showed no spatial structuring, it does not appear that this pattern of IBD is confounded with either an urban/rural cline or any of the human transportation networks. However, the two distant southern sites, both of which were less urbanised and more isolated from the highway network than the northern sites, exhibited higher genetic distances that could be due to any combination of reduced active dispersal leading to IBD, reduced gene flow along highways, or reduced gene flow between urban and rural areas, as has been proposed [25, 46].

Mantel tests on individual genetic distances showed only marginally higher spatial dependence (r = 0.062, P < 0.001) among *Ae. albopictus* separated by tens of kilometres than that observed in a study of an urban *Ae. aegypti* separated by less than 3.5 km (r = 0.047, P < 0.05) [31]. Considering that the flight range of *Ae. albopictus* is only slightly greater than that of *Ae. aegypti* [17, 84], the weaker spatial dependence of structure in *Ae. albopictus* may reflect higher dispersal rates along human transport networks in this mosquito. We found that individuals sampled greater distances from shipping terminals were generally more genetically distinct than those sampled nearer to either network. Guangzhou has recently taken several steps to reaffirm its position as a major regional shipping hub [85], and maintains many local, national and regional shipping routes that could allow for passive mosquito dispersal. However, it is unclear from our results whether the genetic similarities between *Ae. albopictus* collected near the ports are due to the local movement of vessels between ports, or to a common source of imported mosquitoes that are distributed to all shipping terminals.

In recently invaded regions, the movement of *Ae. albopictus* along highways is thought to facilitate dispersal inland from coastal invasion sites [14, 38, 86]. Considering the positions of Jiuwei Village and Zhucun Village (Sites 8 and 9), which had multiple closely related individuals spread between them, transport along highways appears the most reasonable explanation. Comparable observations in *Ae. aegypti* imply a similar dispersal process [31]. However, despite the extensive highway network within Guangzhou, our statistical analyses found that the highway effect on gene flow was weaker than the effect of ports or rivers. This may be in part due to the strong statistical correlation between geographic distance and “Highway Isolation”. In addition, most studies investigating gene flow in *Ae. albopictus* have been conducted in areas much less urbanised than Guangzhou, such as within Madagascar and Réunion [45, 46]. The highway connectivity derived in these studies implies a much less comprehensive highway network than in Guangzhou. The relationship between the highway network and the Breteau Index shows that the most highly connected sites also have the lowest *Ae. albopictus* density. Thus, while highly connected sites may have higher than average migration rates, the total number of migrants along the network will be low (*Nm* [65], analogous to population density * network connectivity). This may help explain why the sample site at the North Campus of Sun Yat-sen University (Site 4) was both the most connected to the highway network and the most genetically distinct. In contrast, Jiuwei Village and Zhucun Village were less connected to the highway network than Site 4 but scored much higher on the Breteau Index.

The findings from this study inform future control projects that rely on releases of sterile individuals (*Wolbachia-*based and others, see review in McGraw and O’Neill [11]) and improve our understanding of the population dynamics of *Ae. albopictus* throughout its range. Releases of *Wolbachia-*infected *Ae. aegypti* have been conducted at several locations around the world by the Eliminate Dengue Program (http://www.eliminatedengue.com/program) and in Kuala Lumpur by *Wolbachia* Malaysia (http://www.nst.com.my/news/2017/03/225454/health-ministry-releases-wolbachia-infected-mosquitoes-keramat). These programs aim to permanently infect local populations with virus-inhibiting *Wolbachia* [87] transinfected from *Drosophila melanogaster* [88]. This differs from the releases undertaken in Guangzhou, in which *Ae. albopictus* males carrying multiple infections of *Wolbachia* are released to mate with wild females, which then produce inviable embryos, leading to local population crash [11]. For either method, reinvasion of controlled regions from external, uncontrolled regions must be prevented to maintain successful control over time. Reinvasion of a target control site was observed in an invaded population of *Ae. aegypti* that exhibited seasonal fluctuations in density, resulting from either active dispersal or human transportation along roads [89]. In Guangzhou, the two release sites were highly connected to either the river network (Dadaosha Island, Site 11) or to both the river and port networks (Shazi Island, Site 12). If suppression releases are undertaken throughout Guangzhou, sites like Shazi Island that are near to shipping terminals may be the most easily reinvaded.

## Conclusions

Our findings point to the complex processes of gene flow among *Ae. albopictus* in Guangzhou, facilitated by human transport networks. Similarly-sized cities could show similar genetic patterns, though this will likely depend on whether the local highway network occupies areas of suitable *Ae. albopictus* habitat. Genetic differentiation may be weaker in recently invaded cities due to founder effects, though this would depend on the frequency of repeated introductions [90] and their populations of origin. Our molecular and analytical approach could be employed to detect inland dispersal between cities, and quantify the relative passive effects of transport (civilian and freight) between bus and railway stations, airports, and truck depots. At global scales, they can be used to assign individuals to regional groups with high confidence [41], allowing for precise modelling of global migration routes throughout the *Ae. albopictus* range.

## Acknowledgments

We would like to thank Yongjun Li and Hongxin Ou for assistance with sample collection and rearing. We also thank the NeCTAR Cloud (https://nectar.org.au/research-cloud/) for providing computational resources.

## Supporting Information

**S1 Table. CDPOP Input File.**

**S2 Table. Results of Mantel Tests on mtDNA Haplotypes.** Mantel tests found no significant relationships between Rousset’s *a* and any of the variables representing barriers or network isolation.

**S1 Fig. Relationship between pairwise mtDNA Nei’s *D* estimates and geographic distance.** Colour codes are the same as in Fig 3. No overall trend of IBD was observed.

## Data Reporting

Demultiplexed *.fastq files have been deposited at NCBI SRA under [name].

Locations of all sequenced individuals are recorded in the file [name.txt].

